# Nextflow vs. plain Bash: Different Approaches to the Parallelisation of SNP Calling from the Whole Genome Sequence Data

**DOI:** 10.1101/2024.02.27.582354

**Authors:** Marek Sztuka, Krzysztof Kotlarz, Magda Mielczarek, Piotr Hajduk, Jakub Liu, Joanna Szyda

## Abstract

This study compared computational approaches to parallelisation of an SNP calling workflow. Data comprised DNA from five Holstein-Friesian cows sequenced with the Illumina platform. The pipeline consisted of quality control, alignment to the reference genome, post-alignment, and SNP calling. Three approaches to parallelisation were compared: (i) a plain Bash script in which a pipeline for each cow was executed as separate processes invoked at the same time, (ii) a Bash script wrapped in a single Nextflow process, and (iii) a Nextflow script with each component of the pipeline defined as a separate process. The results demonstrated that on average, the multi-process Nextflow script performed 15% to 27% faster depending on the number of assigned threads, with the biggest execution time advantage over the plain Bash approach observed with 10 threads. In terms of RAM usage, the most substantial variation was observed for the multi-process Nextflow, for which it increased with the number of assigned threads, while RAM consumption of the other setups did not depend much on the numbers of threads assigned for computations. Due to intermediate and log files generated, disk usage was markedly higher for the multi-process Nextflow than for the plain Bash and for the single-process Nextflow.

## 1. Introduction

In animal genomics, the rapid development of high-throughput technologies during the past few decades has seen a considerable increase in the availability of data (Cao et al 2018, Routhier and Mozziconacci 2022). Among them, the most common data structure is the whole genome sequence (WGS) that is now available to thousands of individuals. For example, the 1000 Bull Genomes Project database for cattle (Hayes and Daetwyler 2019) currently harbours polymorphic variants identified from WGSs of over 5000 individuals. Not only the efficient storage of WGS data and stable processing pipelines are essential for the analysis, but also the whole pipeline from the raw fastq files to Variant Call Format (VCF) files containing identified variants has to be completed in a time-effective manner using systems that are executed in a parallel mode and are robust towards the fluctuation computational resources available at run-time. The so-called rWGS (rapid WGS) is an emerging topic in the analysis of bio-data (see e.g. Sweeney et al 2021), including cattle, for which fast variant identification may have implications on fast selection decisions. Therefore, effective, efficient, and robust computing approaches are becoming increasingly important (Cios et al 2005, Asgari and Mofrad 2015) and so is the pipeline management system software.

There exists a plethora of workflow management systems, ranging from open-source solutions e.g. Jenkins (www.jenkins.io) or Snakemake (Mölder et al 2021) to commercial software such as the Automic Automation (www.broadcom.com). Moreover, platform-dedicated systems are also available, like the AWS Code Pipeline for users of Amazon Web Services. The Nextflow pipeline management system (Di Tommaso et al 2017) has recently gained popularity, mainly within the field of genomics and, more broadly, bioinformatics, which is to a large extent due to its simplicity of implementation, good tutorials, community support, and most importantly thanks to the availability of built-in directives dedicated to standard processing of WGS pipelines that are missing in general-purpose pipeline management software. One of the important aspects of using the Nextflow system is the automatic parallelisation and scaling of data processing from local computers to clusters, both within and across individual WGS samples, which accelerates the execution of computationally intensive tasks. It also enables the use of multiple scripting languages, including Bash, R, and Python, which are very popular within the bioinformatics community.

Our study aimed to compare the computational efficiency and hardware requirements of the native Bash implementation with the implementation through the pipeline management system. The Nextflow DSL2 (domain-specific language) pipeline management system and the context of detecting single nucleotide polymorphisms (SNPs) in the WGS data were chosen as an example management system and a pipeline, respectively. Since all the elements of the pipeline are required for obtaining the final outcome – the VCF of called SNP genotypes. The practical aspect of the underlying comparison was to present the overall runtime of the entire workflow, without splitting between memory, time, and disk usage of particular stages.

## 2. Material and methods

### 2.1. Animals and DNA-sequencing

The genomic DNA of five Holstein-Friesian cows was sequenced with the Illumina HiSeq2000 platform in the paired-end read mode with a read length of 100 bp. The number of reads available for a single animal ranged between 391952216 and 407377182. In this study, for demonstration purposes, only sequence reads mapped to chromosome 25 (BTA25) were processed.

### 2.2. Bioinformatic analysis

The bioinformatics pipeline for SNP calling consisted of: (1) the quality control step performed using fastQC software to assess the quality of the raw DNA sequence reads, (2) the alignment of sequence reads to BTA25 from the ARS-UCD1.2 reference genome (NCBI accession number: PRJNA391427) with BWA-MEM software (Li and Durbin 2009), (3) the post-alignment processing, and (4) SNP calling with Samtools package (Li et al 2009).

The pipeline defined above was executed using three different setups visualised in Figure 1: (1) as a plain Bash script run in parallel for each of the five individuals (**plain Bash)**, (2) as an entire Bash script wrapped into a single Nextflow process (**single-process Nextflow)**, (3) as each component of the Bash script, corresponding to each step of the SNP calling pipeline, defined as a separate Nextflow process, with processes connected via channels (**multi-process Nextflow**). Each setup (1-3) was executed in a parallel mode across each cow, additionally with multiple numbers of threads defined within each cow. The following constellations of the numbers of threads (T) and the numbers of forks (F) represented by a cow-level process were implemented: F5T1, F5T5, F5T10, and F5T15. In the **plain Bash** setup, parallelisation across individuals was implemented by executing the full pipeline for each cow as a separate process. In **single-process** and **multi-process Nextflow** setups, Nextflow was used to implement across-cow parallelisation. Furthermore, to compare the parallelisation strategies implemented via Nextflow, the **multi-process** pipeline was executed with 50 threads, but sequentially processing each cow (F1T50).

**Fig. 1.**
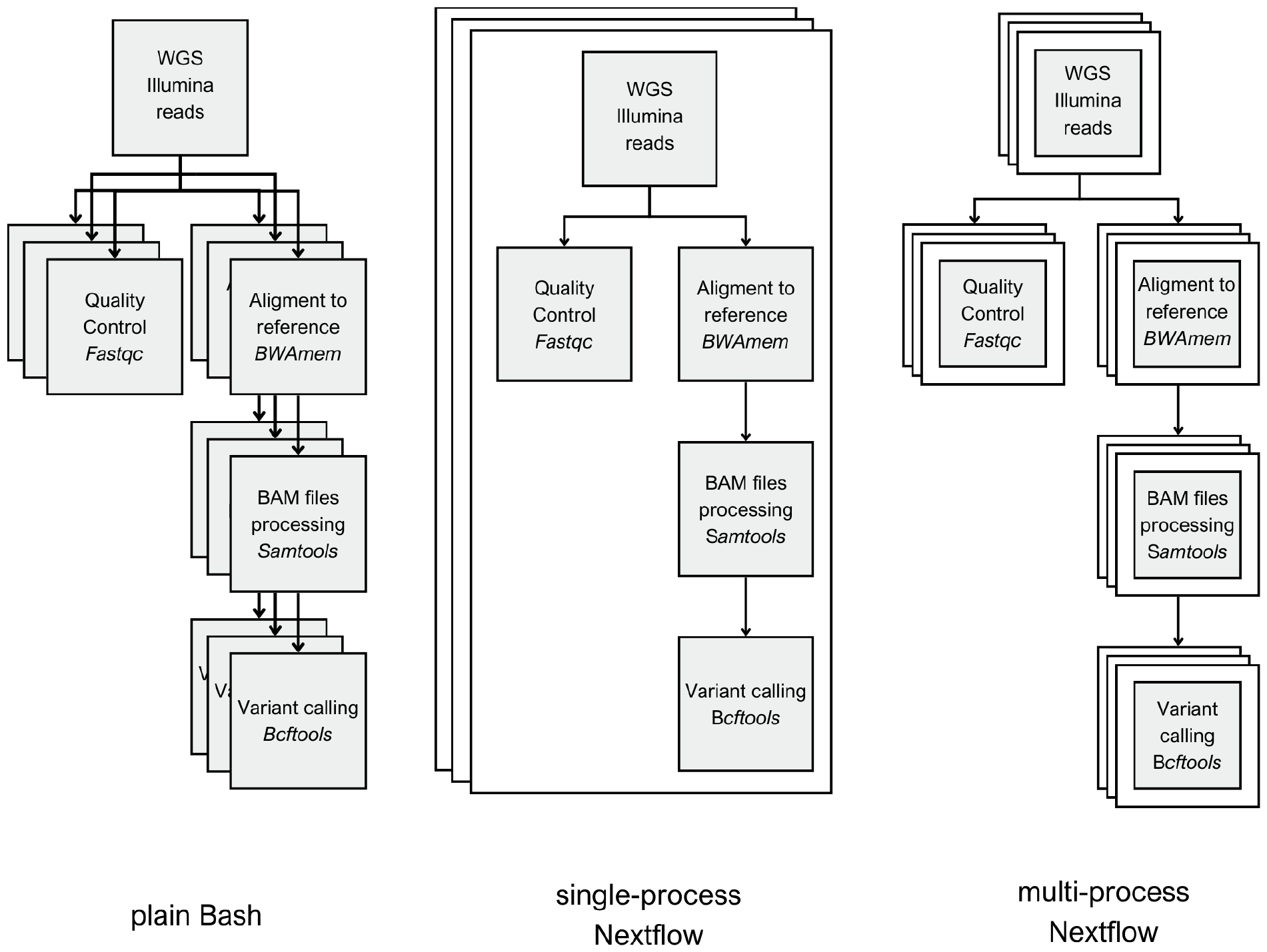
SNP calling pipelines implemented in the study. From left to right, (1) not managed, plain Bash, (2) single-process Nextflow where the entirety of the pipeline was crammed into one Nextflow process, (3) proper Nextflow pipeline design in which each step of the pipeline was a separate Nextflow process, connected via channels.

All setups were compared for execution time, maximum memory usage, and hard disk storage space.

### 2.3. Hardware

All computations were performed on a server equipped with two Intel Xeon CPUs (E5-2699 v4) with 44 threads each, with a base frequency of 2.20 GHz and 188 GB of RAM. During the execution of the pipelines, the server was dedicated solely to our analysis. Only housekeeping processes were active along with pipeline executions.

## 3. Results

The outcome of the plain pipeline consisted of five VCF files generated for BTA25 in the single-individual mode, that is, separately for each cow for which the number of SNPs varied between 312100 and 353855. Outputs comprised quality reports on sequenced reads from fastQC software in HTML format, as well as text log files. Additionally, for pipelines implementing Nextflow, reports in HTML format with information on pipeline execution were generated. Each execution setup (**plain Bash, single-process Nextflow, multi-process Nextflow**) resulted in the same set of identified SNPs.

Regarding execution time (Figure 2), **multi-process Nextflow** was the fastest regardless of the number of threads assigned, except for sequential implementation on only one core (F5T1), where **plain Bash** was the most computationally efficient, being 13.80% faster than **multi-process Nextflow** and 19.80% faster than **single-process Nextflow**. On the contrary, for the parallelised computations, the advantage in execution time of **multi-process Nextflow** over **plain Bash** varied between 15.71% and 21.15% and between 23.10% and 26.79% over **single-process Nextflow**, depending on thread configuration. The largest difference was observed for F5T10, when **multi-process Nextflow** executed 11 hours and 25 minutes, while **plain Bash** ran for 15 hours and 30 minutes. Interestingly, no marked differences in execution time were observed between setups of 10 and 15 cores per animal (approximately 20 minutes). When comparing **plain Bash** with **single-process Nextflow**, there was no clear winner in terms of execution times, since they were very similar. Regardless of the implementation, the largest decrease in execution time occurred when the processing of each animal was assigned 5 cores (F5T5), compared to serial execution F5T1. The F5T5 configuration was almost 6.5 times faster than the serial implementation.

**Fig. 2.**
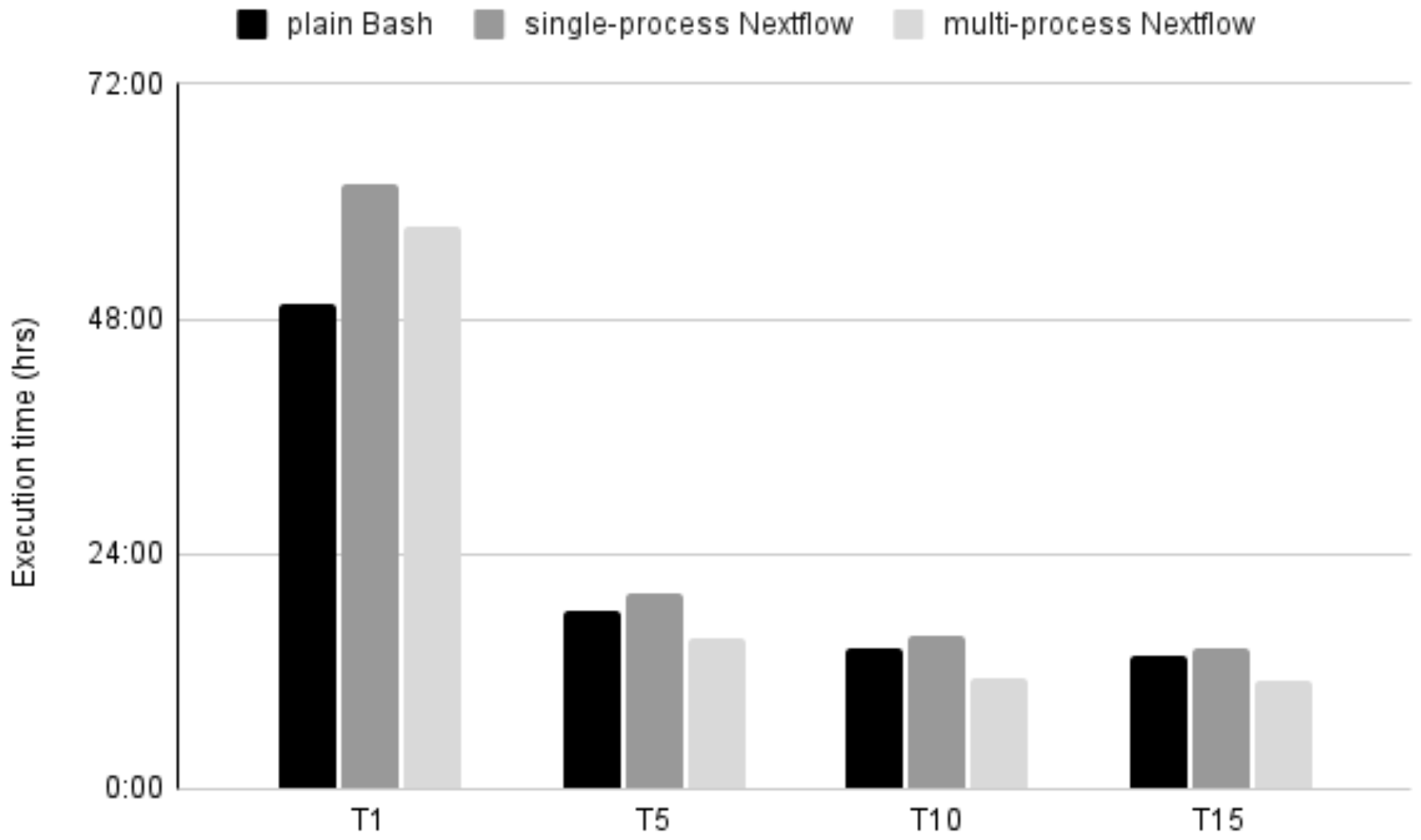
Average execution times of various setups of the SNP calling pipeline. On the X axis there are a number of threads used per individual. Differences in computation time between animals were not significant.

When comparing parallelisation implemented via Nextflow with internal parallelisation implemented in the programmes that are the components of the pipeline, i.e. fastQC, BWA-MEM, and Samtools, the advantage of splitting the entire pipeline into separate processes corresponding to each individual (F5) became evident (Figure 3). F5T10 **multi-process Nextflow** configuration separately assigns 10 threads to each of the 5 cows executed in 11 hours and 29 minutes, while the sequential approach computing one cow after another with 50 threads assigned for each animal (F1T50), despite the same resources defined, ran over three times longer (34 hours and 14 minutes).

**Fig. 3.**
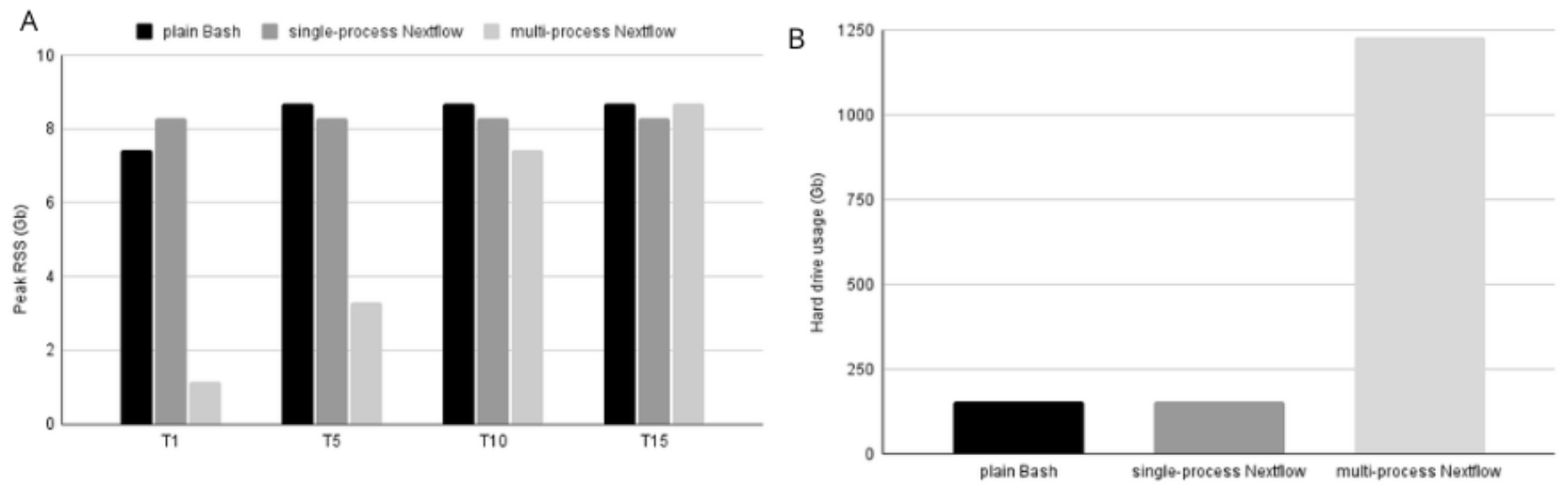
A. Maximum RAM utilisation of various setups of the SNP calling pipeline. B. Usage of hard drive by different implementations of the SNP calling pipeline.

Regarding RAM utilisation (Figure 4A), a large difference was observed between **multi-process Nextflow** and the other two setups in all thread constellations implemented. In multi-process Nextflow, RAM consumption increased with the number of assigned threads, while in the other setups, it did not depend much on the numbers of threads assigned for computations. This result clearly demonstrates the superiority of Nextflow in memory management. The largest difference between the three setups appeared for the F5T1 constellation, for which the **multi-process Nextflow** pipeline used 1.16 GB of memory, while the other setups consumed up to 7.44 GB (**plain Bash**) and 8.26 GB (**single-process Nextflow**). Therefore, the **multi-process Nextflow** was 7 times more memory efficient. In terms of hard drive space requirements (Figure 4b), **multi-process Nextflow** occupied a markedly larger disk space, 1227 GB, compared to 156 GB used by **single-process Nextflow** and the **plain Bash** script. These results were consistent in all configurations. This disparity is primarily due to the creation of a working directory in which Nextflow generated temporary files were stored.

## 4. Discussion

With the rapidly growing popularity of the Nextflow workflow management system, it is important to implement the available tools in the most effective way to maximise profit in execution time and computer resources (Bielecki and Śmiałek 2023). Published reports of genomic workflow comparisons are scarce and do not formally compare the same pipeline implemented with and without a workflow management system. Recently, (Hu et al 2022) proposed a Nextflow pipeline for single-cell ATAC-seq data analysis and compared it with two other pipelines implemented without workflow management system (https://github.com/wbaopaul/scATAC-pro) and with the Snakemake workflow management (https://github.com/liulab-dfci/MAESTRO). Interestingly, all three pipelines produced different results, but with regard to memory consumption and execution time, no marked differences between implementations emerged. The authors of this study stated that the Nextflow pipeline was characterised by a much higher level of flexibility and ease of parameter optimisation. Mpangase et al. (2021) created a Nextflow pipeline for obtaining raw read counts from RNA-seq data and compared it with the Rsubread package (10.1093/nar/gkz114) implementing the pipeline in R. However, since both implementations used different software, observed differences in execution times or memory usage are not meaningful in the context of a comparison of workflow efficiency.

An added value of using Nextflow pipeline management systems is the presence of the nf-core library that provides a platform where researchers can contribute and share their analysis workflows that even aim to become standardised workflows for processing various types of *omic* data (Ewels et al 2020). Furthermore, the platform also provides good documentation that facilitates pipeline implementation. Workflow management systems address the problem of pipeline portability and reproducibility, which pose a serious problem in many research areas (Grüning et al 2018, Kim et al 2018). Moreover, the managed system application provides visualisation tools during the execution of processes and after their completion, helping to compare and tune pipeline execution parameters. Another very practical feature is the ability to resume a process after a halt. This negates the need to run the process from the very beginning after resolving a problem, which makes the debugging process more efficient. The downside of this feature, however, lies in creating a work directory that consumes substantial amounts of drive space. Nextflow also enables the user to specify the number of threads used on several forks for each process of the pipeline.

The feature of using a pipeline management system is sharing common data across processes that does not enforce repeated computations when it is not necessary, e.g. genome indexing. Our comparison demonstrated that, for the shared memory architecture used for computations, Nextflow workflows are more efficient in terms of memory and CPU management of processes running in parallel. The most important benefit is related to the fact that Nextflow implements the functional reactive programming paradigm that supports non-synchronous data processing through defining so-called channels that transfer data between parallel processes, so that, in practice, when computations are completed for one animal, available resources are reallocated to other animals. Separating tasks into channels also allows for running the quality control independently of the alignment process, which is not possible under **plain Bash** implementation. However, it should be realised that as the number of defined threads per cow increases, **multi-process Nextflow** begins to use more of the available computing power, resulting in higher memory usage. Still, from a certain point on, increasing the number of threads did not result in a marked benefit in terms of execution time. Although the aspect was not formally investigated in our study, we suspect that the increased computational load of handling multiple parallel processes impeded the benefits of parallel computations, especially in the pipeline (like ours) that contains components that do not execute in the parallel mode, or do not employ parallel processing for the majority of its computations, e.g. the Samtools package, which uses multithreading exclusively for compressing alignment map files but not for their downstream processing. Although parallel computations provide important benefits for the overall execution time, this benefit is hampered by overhead of multithreading that is mainly composed of thread management, context-switch costs, and cash repletion (see already e.g. Kwak et al 1999).

A missing aspect of our study was the comparison of workflow performance implemented in a distributed memory architecture. Although due to the lack of the appropriate computing environment dedicated entirely to the comparison, i.e. running no other processes, it can be speculated that **multi-process Nextflow** implementation would be even more beneficial over **single-process Nextflow**. Furthermore, the use of the **plain Bash** approach would require manual implementation of the MPI directives, which would impose an additional programming burden. It is also worth mentioning that HDD access is a critical point of pipeline runtime. The benefit of using a management system is that it handles HD IO operations within its processes that allows to optimize resources management, including HDD IO.

## 5. Conclusions

In dairy cattle, we currently experience fast-growing load of digital data that on the phenotypic and environmental level originates from precision livestock farming systems utilised on many farms as well as on the genomic level – originating from sequencing of whole genomes of many individuals, mainly bulls. The expectation is that this information will be routinely used in dairy management and breeding decisions. In view of those fast-growing sizes of data, workflows and code parallelisation are very important computational aspects. In this context, the Nextflow workflow management system is a useful tool not only for managing pipelines, which strongly and efficiently supports computational parallelisation and enables the user to specify the number of threads used. Still, it is important to consider that in parallel computing, a critical element is the proper design of the computing architecture expressed by the number of computing tasks and available CPU cores (Akon et al 2005). Efficient resource management guarantees that these multiple tasks coexist without interfering with one another, resulting in optimal system performance, so that a well-designed system implementing optimal number of threads and a number of parallel computing individuals leads to optimal computational performance.

## Acknowledgements

-

## Availability of data and materials

The DNA sequences of the 5 cows are available from the NCBI BioProject database under the accession ID: PRJNA359667 (Szyda et al 2015). Accession numbers corresponding to particular samples are as follows: (HOLPOL2 - SRX2455298, HOLPOL3 - SRX2455321, HOLPOL23 - [SRX2455312](https://www.ncbi.nlm.nih.gov/sra/SRX2455321%5Baccn%5D), HOLPOL25 - SRX2455300, HOLPOL27 - SRX2455319). The code is available on the GitHub repository: https://github.com/paq88/Variant_Calling_Nextflow_pipeline

